# PERSIST: A programmable RNA regulation platform using CRISPR endoRNases

**DOI:** 10.1101/2019.12.15.867150

**Authors:** Breanna DiAndreth, Noreen Wauford, Eileen Hu, Sebastian Palacios, Ron Weiss

## Abstract

Regulation of transgene expression is becoming an integral component of gene therapies, cell therapies and biomanufacturing. However, transcription factor-based regulation upon which the majority of such applications are based suffers from complications such as epigenetic silencing, which limits the longevity and reliability of these efforts. Genetically engineered mammalian cells used for cell therapies and biomanufacturing as well as newer RNA-based gene therapies would benefit from post-transcriptional methods of gene regulation, but few such platforms exist that enable sophisticated programming of cell behavior. Here we engineer the 5’ and 3’ untranslated regions of transcripts to enable robust and composable RNA-level regulation through transcript cleavage and, in particular, create modular RNA-level OFF- and ON-switch motifs. We show that genomically introduced transgenes exhibit resistance to silencing when regulated using this platform compared to those that are transcriptionally-regulated. We adapt nine CRISPR-specific endoRNases as RNA-level “activators” and “repressors” and show that these can be easily layered and composed to reconstruct genetic programming topologies previously achieved with transcription factor-based regulation including cascades, all 16 two-input Boolean logic functions, positive feedback, a feed-forward loop and a putative bistable toggle switch. The orthogonal, modular and composable nature of this platform as well as the ease with which robust and predictable gene circuits are constructed holds promise for their application in gene and cell therapies.

## INTRODUCTION

Transcription factor-based gene regulation in mammalian systems remains a cornerstone of both basic biological research and the field of synthetic biology. Transcription factor-promoter pairs such as the Tet-On or Tet-Off systems [30, 61, 74] have been used extensively for both studying genes of interest in research settings as well as controlling transgenes for biotechnology applications. The robust behavior, modularity, composability and large design space of transcriptional regulators has also enabled their layering into multi-transcription-factor circuits for the forward-design of complex cellular behavior. However, hurdles such as epigenetic silencing challenge the therapeutic potential of these strategies.

Epigenetic silencing of transgenes has been observed in many therapeutically-relevant cell types including stem cells [50], neurons [75], and CHO cells [40] and inhibits the engineered functions of these cells over time. While the location of genomic integration has been shown to be one factor that effects silencing [43, 36, 58, 52, 55, 53], other emerging factors such as features of promoter sequences [35, 40, 70, 51, 68, 4] and preferences for a transcriptionally active state [6, 15, 22, 50, 73] make transcription-factor based gene regulation unsuitable in many contexts.

Efforts have turned to developing post-transcriptional platforms comprising activator- and repressor-like regulators. Such platforms would allow the use of constitutive promoters routinely used in gene therapy, which tend to resist silencing [62, 54], and may also find use in modern therapeutic modalities such as mRNA gene therapies. Recently-developed protein-regulation platforms using orthogonal proteases [14, 27, 21], have begun to address this need by controlling protein degradation. The field would benefit from development of protein-based RNA-regulation platforms with similar properties that enable control of protein production, e.g. for regulation of secreted proteins. Some protein-based RNA-level regulators exist, mainly using translational repressors [56, 60, 66] and RNA-cleaving Cas proteins [10, 1, 2, 16, 59, 41, 69], but there is still a great need for demonstrated scalability and composability. Also, all of these examples canonically act as only repressor-like “OFF” switches, where a few RNA-based activator-like “ON” switches have been developed [9, 10, 18, 56, 3], none of which are composable into circuits. The field currently lacks an RNA-acting activation/repression toolkit that is scalable, robust, modular and composable to replace transcription factors.

Here we develop RNA-level ON-switch and OFF-switch motifs that can be engineered into the untranslated regions (UTRs) of transcripts to allow for post-transcriptional gene regulation that is modulated by RNA cleavage. This regulatory modality is less vulnerable to silencing compared to transcriptional regulation because it can make use of vetted constitutive promoters routinely used in gene and cell therapies. The ON and OFF switches can be modified to respond to a variety of RNA cleavage effectors including endonucleases, miRNAs and ribozymes. We term this platform “Programmable Endonucleolytic Scission-Induced Stability Tuning” (PERSIST). In particular, we identify CRISPR-specific endoRNases as useful effectors for RNA cleavage-based regulation and select a set of nine that are orthogonal in their cleavage to serve as RNA-level “activators” and “repressors”. These endoRNases exhibit up to 300-fold dynamic range as repressors and up to 50-fold dynamic range as activators in the PERSIST platform. We demonstrate that our platform exhibits modularity and composability by creating multi-level cascades, all 16 two-input Boolean logic functions, and a positive feedback motif. An interesting feature of these endoRNase regulators is that they can function as both activators and repressors simultaneously, which allows for compact engineering of previously complex motifs such as a feed-forward loop and RNA-level bistable switch. The performance and simplicity of the PERSIST platform, combined with its resistance to silencing, will extend the functional lifetime of more useful engineered genetic circuits for applications in biomanufacturing, gene therapies, and engineered cell therapies.

## RESULTS

### Development of RNA-level switches that respond to RNA cleavage and resist epigenetic silencing

We first set out to develop RNA-level switches that enable turning transgene expression on or off through regulation of transcript degradation. Previous studies have shown that transcript cleavage, for example by miRNA [25] or endoribonucleases [10, 49, 72], can reduce transgene expression (Figure 1a, left). Our initial object was to develop an RNA-level ON switch that activates gene expression in response to transcript cleavage. We envisioned this motif to have three domains (see Figure 1a, right): (1) degradation signals that cause the rapid degradation of the transcript, (2) a cleavage domain which allows the removal of the degradation tag, and (3) a stabilizer that allows efficient translation and protects the mRNA after the removal of the degradation signal. Thus, the transcript is degraded in the absence of a cleavage event and stabilized post-cleavage.

**Figure 1.**
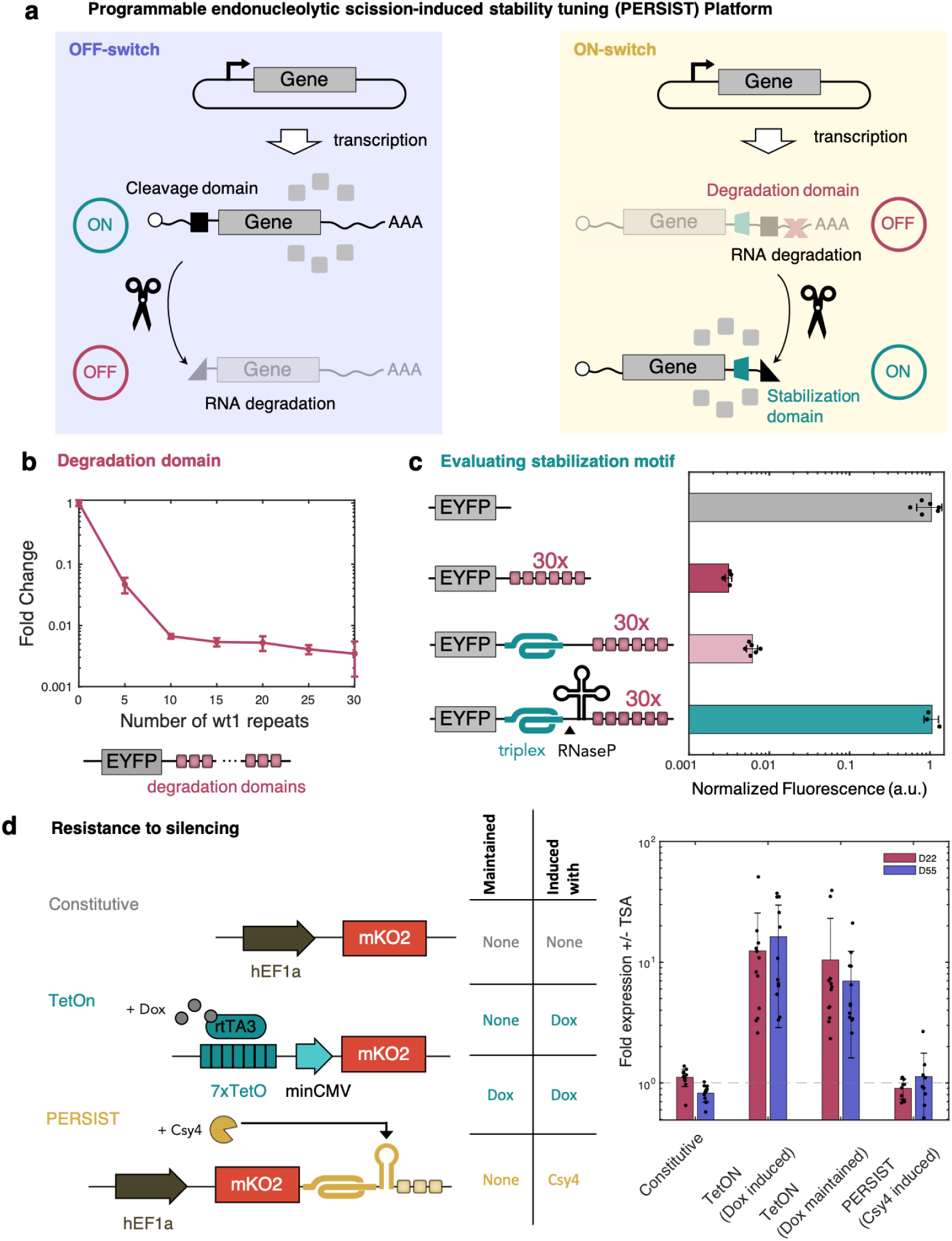
Engineering PERSIST OFF- and ON- motifs for regulated gene expression. **a**, Schematic of RNA-based OFF-switch and ON-switch motif designs that are regulated by RNA cleavage. **b**, Gene expression reduction due to appending increasing numbers of degradation motifs. **c**, Triplex structure rescues gene expression only after removal of degradation motifs by RNaseP. **b,c** n≥2 biologically independent samples where each sample represents the evaluation of > 1,000 transfected cells (HEK293FT). Data are presented as mean ± s.d. **d**, PERSIST shows resistance to epigenetic silencing in clonal populations. Three different constructs (left) each expressing mKO2 under control of a different switch (constitutive, Tet-On or PERSIST-ON) were genomically integrated and single-cell sorted to evaluate clonal silencing effects. The table (middle) shows the maintenance and induction conditions of each cell line type. 22 and 55 days after sorting, 12 clonal lines for each switch were induced with either 4uM Dox or transfected with Csy4 both in the presence and absence of TSA. The ratio of +/- TSA for mKO2 expression was calculated for each cell line in each sample type where a ratio > 1 indicates rescue of response by TSA. While the Tet-On system shows increased mKO2 expression in the presence of TSA which indicates epigenetic silencing effects, the PERSIST switch resists silencing to similar levels as the constitutively expressed reporter. n≥10 independently sorted and evaluated samples where each sample represent the evaluation of ≥ 5 CHO-K1 cells. Data are presented as mean ± s.d.

First we sought to identify a mechanism that would lead to rapid mRNA degradation. A natural short 8bp RNA motif has been recently identified that binds heterogeneous nuclear ribonucleoproteins, which in turn recruit deadenylase complexes, resulting in mRNA degradation in a variety of cell types [28]. We created EYFP reporters that contain varied numbers of repeats (up to 30) of the degradation motif, “wt1”, within the 3’ UTR (Supplementary Figures 1 and 2) and evaluated expression via transient transfection into HEK293FT cells. A reporter containing 30 repeats was efficiently degraded with over 300 fold reduction in fluorescence, while fewer numbers of repeats allowed for varied degrees of repression (Figure 1b, Supplementary Figure 3). Notably, while various methods exist to program RNA production, protein production and protein degradation (e.g. with promoter design [32, 20], upstream ORFS [19], and destabilization domains [7, 8]) respectively), these RNA degradation motifs represent a new way to fine-tune mRNA half-life.

Next, we searched for a sequence that would stabilize the transcript and allow for translation after removal of these degradation tags. Importantly, this element should not stabilize the transcript when the degradation tags are still present. An RNA triple helix (triplex) structure from the MALAT1 long noncoding RNA [65, 11, 71] has been used in several synthetic biology studies to stabilize engineered transcripts [49, 46, 47]. When we inserted the MALAT1 triplex upstream of 30 repeats of the wt1 degradation motif, expression of the EYFP reporter remained low (Figure 1c). When we included a mascRNA sequence, which is cleaved naturally by RNase P, between the triplex and degradation motif, expression was restored to constitutive levels (Figure 1c). Overall this ON-switch exhibits a dynamic range of 166-fold, which is promising for its adaptation as an RNA ON-switch motif for transgene regulation.

After the creation of the RNA ON-switch, it was important to evaluate its ability to resist epigenetic silencing. We first developed an endoRNase-responsive ON switch using Csy4. The CRISPR endoRNase Csy4 (PaeCas6f) has been used as a translational repressor in mammalian synthetic biology previously [10, 49, 72] and here we show that it can be repurposed to activate expression when its cleavage site is placed within the PERSIST-ON motif (Supplementary Figure 4). We chose to benchmark the Csy4-responsive PERSIST ON-switch against the routinely-used Tet-On system [31]. In the Tet-On transcriptional ON-switch, rtTA3 is induced by doxycycline (Dox) to activate expression at the TRE promoter. We engineered both the Tet-On and PERSIST switches to drive expression of the fluorescent protein mKO2 and compared to a control construct encoding constitutive mKO2 expression from a hEF1a promoter (Figure 1d). All three constructs were integrated into landing pads located in the putative Rosa26 locus of CHO-K1 cells [24] and polyclonal populations showed expected responses (Supplementary Figure 5a). We then used single cell sorting to generate 12 cell lines for each of four different conditions tested: (1) constitutive expression, (2) Tet-On switch maintained without Dox, (3) Tet-On switch always maintained in the presence of Dox to simulate constitutive expression and (4) PERIST ON-switch (Supplementary Figure 5b). To evaluate response, each of the switches were induced (either by Dox addition or Csy4 transfection) 22 days and 55 days after sorting. We measured mKO2 levels in the presence and absence of the histone deacetylase Trichostatin A (TSA) (Supplementary Figure 5c), which would rescue any loss in response specifically due to silencing. As seen in Figure 1d both the constitutive reporter and the PERSIST ON-switch respond similarly with and without TSA as indicated by their +/-TSA ratio being close to one. This would imply that both of these constructs resist epigenetic silencing. Interestingly, rescue with TSA addition was observed for the Tet-On system regardless of whether or not it was maintained in Dox, implying that both cases experience epigenetic silencing. Whether this effect is attributed to properties of the Tet-On promoter sequence itself or the transcriptional activator, it is clear that the PERSIST platform avoids these pitfalls and enables long-term robust yet regulatable response. The reliability of the response suggests that our post-transcriptional platform for transgene regulation could prove useful for applications where long-term control is required such as in cell therapies and biomanufacturing.

### The PERSIST platform responds to CRISPR endoRNases, ribozymes and miRNAs

In order to generate a library of RNA-regulators, we next sought to evaluate whether other endoRNases besides Csy4 could cleave the transcript and activate the ON-switch motif or repress the OFF-switch motif. The Cas6 family that Csy4 belongs to and the Cas13 family of CRISPR-specific endonucleases cleave pre-crRNAs to produce shorter gRNA sequences used for DNA- or RNA-targeting respectively [34]. Importantly, these endonucleases cleave short, specific (often hairpin-structured) sequences called direct repeats, and orthologs of these endonucleases are thought to cleave their respective direct repeats orthogonally [48]. We hypothesized that these Cas6 and Cas13 families could provide a large number of parts that orthogonally cleave mRNAs when their cognate cleavage domain is engineered into the transcript. We evaluated 9 total endonucleases, four from the Cas6 family: Csy4, Cse3 (TthCas6e)[57], CasE (EcoCas6e)[38] and Cas6 (PfuCas6-1)[13] and five from the Cas13 family: LwaCas13a [1], PspCas13b [16, 59], PguCas13b [16, 59], RanCas13b [16, 59], and RfxCas13d [42]. Since Csy4 and all of the Cas13 family endonucleases have previously been used in mammalian cells, only Cse3, CasE, and Cas6 required mammalian codon optimization. Several other modifications to the Cas13 family endonucleases were made to improve performance and ensure compatibility with other endonucleases, such as removing previously-required reporter-fusions and including nuclear localization signals (data not shown). We observed robust ON responses to most of these endoRNases when their cognate recognition sequences were placed between the triplex and degradation signal in the 3’UTR-located PERSIST-ON-switch motif (Figure 2a and Supplementary Fig 4 and 6). Interestingly, for some Cas endoRNases, the triplex was not required for ON-switch functionality (Supplementary Fig 6), likely due to the fact that the endoRNases stay bound to their cleaved product and protect the 3’ end of the transcript as previously shown [33, 44, 12]. These same set of endonucleases led to robust OFF responses of up to 300-fold repression when their target sequences were placed in the transcript outside of the PERSIST ON-switch with larger repression occurring when placed in the 5’ UTR of the transcript for those tested (Figure 2a, Supplementary Figure 7). We also verified that the PERSIST ON-switch responded to the endoRNase CasE in several other relevant cell lines: CHO-K1, HeLa, and Jurkat (Supplementary Figure 8), suggesting that the native degradation and stabilization mechanisms by which our PERSIST platform functions are ubiquitous. Our optimization provided a final set of 9 endoRNases that act as repressors when their recognition sites are placed in the 5’ UTR of transcripts (PERIST OFF-switch) and as activators when their recognition sites are placed in the the 3’ UTR PERSIST ON-switch (Figure 2a).

**Figure 2.**
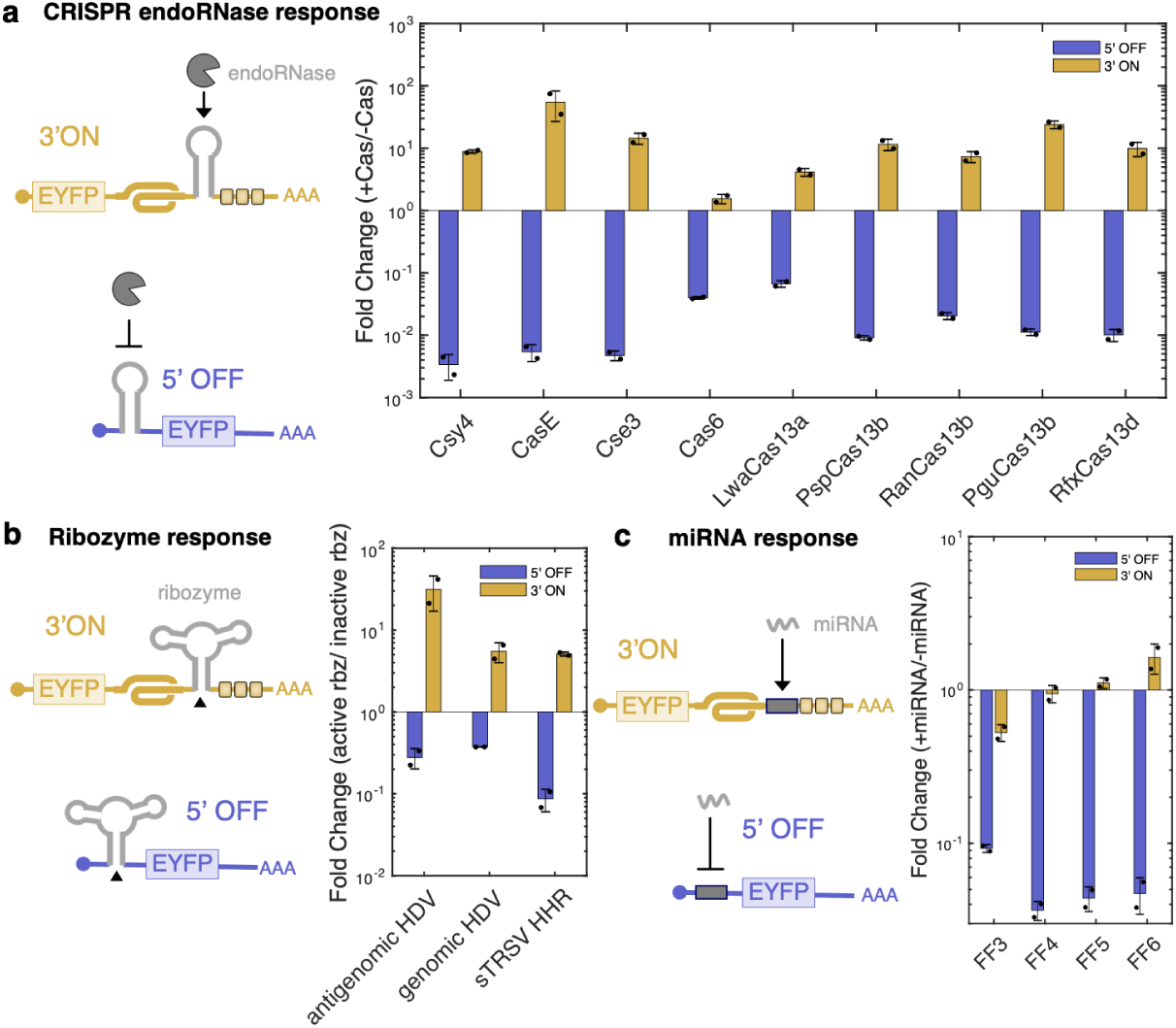
Evaluating the PERSIST platform with various cleavage effectors. **a**, CRISPR endoRNases activate the PERSIST-ON motif and repress the PERSIST-OFF motif. **b**, Ribozymes activate the PERSIST-ON motif and repress the PERSIST-OFF motif. **c**, Some miRNAs activate the PERSIST-ON motif and all repress the PERSIST-OFF motif. **a,b,c** n≥2 biologically independent samples where each sample represents the evaluation of > 1,000 transfected cells (HEK293FT). Data are presented as mean ± s.d.

In addition to endoRNases, we also evaluated whether the PERSIST platform could be adapted to respond to other RNA cleavage events. Ribozymes placed in the PERSIST-OFF and PERSIST-ON motifs were able to repress and activate gene expression respectively as compared to their cognate inactive mutant versions (Figure 2b). This could be useful for detecting endogeneous ribozymes or regulating expression based on small molecules via aptazymes. We were also especially interested in adapting the PERSIST platform to respond to miRNAs. Typically, reporters that detect high levels of miRNA and provide a positive readout (i.e. “miRNA high sensors”) have been difficult to design and optimize in part because they often require a double-inversion step [67] or multiple parts [63]. We modified the PERSIST-ON switch motif to contain a miRNA target site between the triplex and degradation motifs and evaluated response to several synthetic miRNAs. Albeit modest in performance, PERSIST-ON miRNA reporter expression increased in response to the synthetic miRNAs FF5 and FF6, while placing miRNA target sites in the 5’ UTR OFF position canonically decreased expression as has been shown previously [25] (Figure 2c)

Overall, the PERSIST switch platform encodes the ability to respond both positively and negatively to RNA cleavage events based on recognition site position. The dynamic range, robustness and adaptability for different types of inputs makes it promising for use in synthetic biology and other genetic manipulation applications.

### PERSIST switches demonstrate orthogonality, modularity and composability

The repression and activation responses to CRISPR endoRNases in the 5’ UTR OFF-switch and 3’ UTR ON-switch positions respectively suggested that these proteins could be used as RNA-level activators and repressors for programmable gene circuits. In fact, the recognition sites function similarly to transcription factor operons, except that the repressor or activator functionality of the endoRNases is determined by the location of their cognate recognition sites in the transcript rather than encoded in the protein itself (Figure 3a).

**Figure 3.**
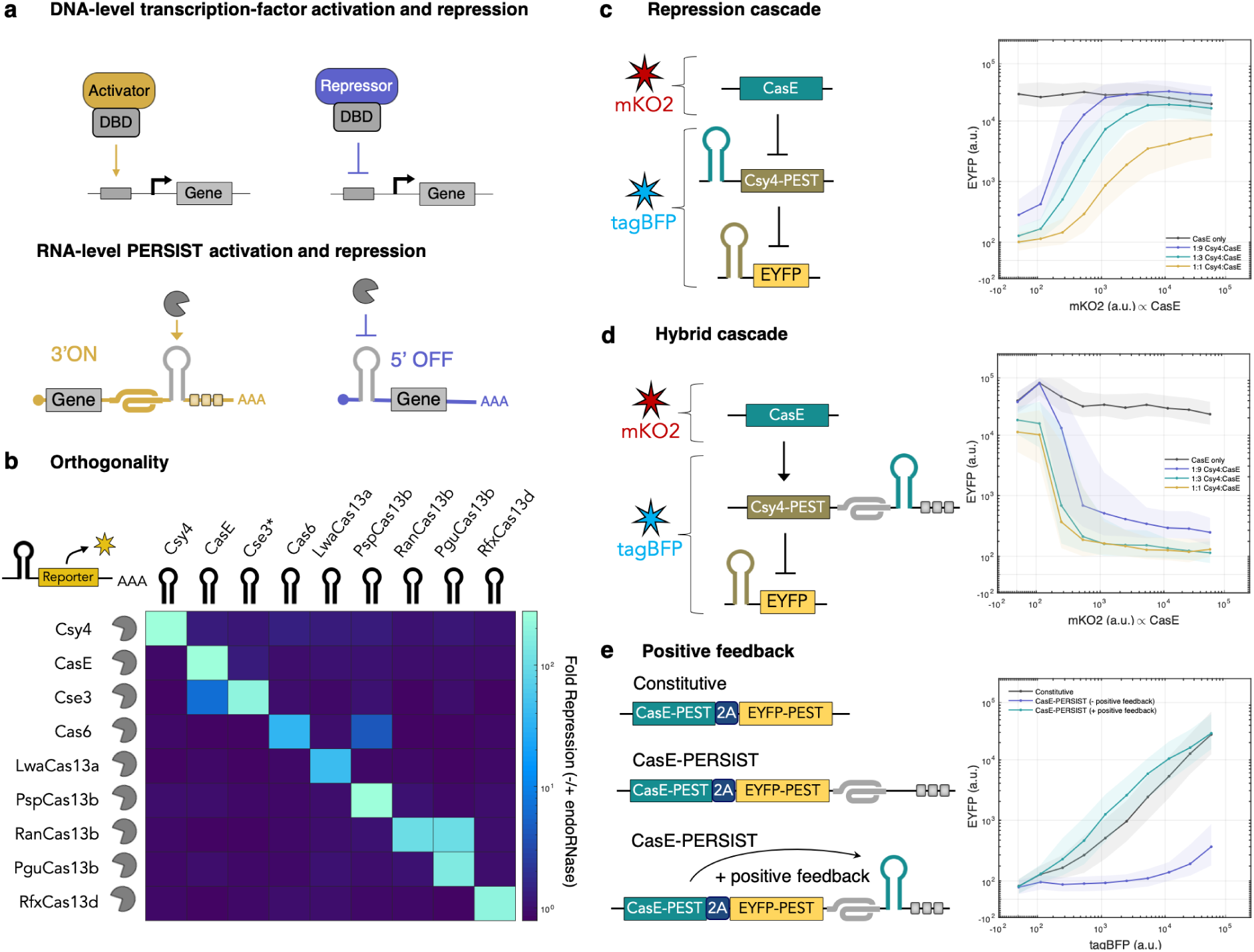
PERSIST endoRNase platform is orthogonal and composable. **a**, Schematic comparing traditional transcriptional programming to post-transcriptional programming with the PERSIST platform. While repressor/activator functions are encoded in transcriptional regulators, the repressive/activating functions of Cas endoRNases are encoded in their programmed cleavage location. **b**, Evaluation of Cas endoRNase pairwise orthogonality. The ability of each Cas endoRNase to repress reporters containing each endoRNase recognition site was evaluated by poly-transfection. Fold repression values are indicated by colorbar. While some pairs should be avoided, most endoRNases cleave orthogonally. **c**, Cas endoRNase repression cascade. An endoRNase can repress another endoRNase repressor through a 2-stage cascade. For plasmid ratio calculations, CasE plasmid was maintained at 75ng. **d**, Cas endoRNase hybrid cascade. An endoRNase can activate another endoRNase repressor through a 2-stage cascade. **c,d**, For plasmid ratio calculations, CasE plasmid was maintained at 75ng. CasE plasmid transfection efficiency was tracked by a plasmid encoding constitutive mKO2 (x-axis) while Csy4 and EYFP reporter plasmid transfection efficiencies were tracked by a plasmid encoding constitutive tagBFP. Median EYFP fluorescence (line) and interquartile range (shaded regions) were evaluated across various mKO2 levels for tagBFP-positive cells. **e**, Cas endoRNase auto positive feedback. An endoRNase can activate itself through a positive feedback loop. We compared a CasE construct tagged with EYFP alone (gray line), a construct containing the PERSIST ON motif but no endoRNase target site (blue line) and a construct containing a CasE target site in the PERSIST motif (green line). Construct transfection efficiencies were tracked by a plasmid encoding constitutive tagBFP. Median EYFP fluorescence (line) and interquartile range (shaded regions) were evaluated across various tagBFP levels. **b,c,d**, Each sample represents the evaluation of > 1,000 transfected cells (HEK293FT).

To demonstrate the utility of our platform, we first sought to evaluate the orthogonality, modularity and composability of the Cas-based PERSIST-switches. In particular, in order to be a platform that could be used in place of transcription-factor-based gene regulation, the endoRNase platform should have several characteristics: (1) the Cas proteins should be orthogonal to each other, i.e. have minimal cross-talk with each others’ recognition sites, (2) the Cas protein-recognition site pairs should be modular such that recognition sites can be placed in either the 5’-OFF or 3’-ON PERSIST switch locations (in any order or combination) and have predictable behavior, and (3) the regulation enabled by the endoRNases should be composable, that is, it should be possible to connect endoRNases to create layered circuits.

#### Evaluating orthogonality

We first evaluated the orthogonality of the Cas proteins by testing each Cas protein with every pairwise combination of Cas-responsive PERSIST-OFF reporters. Notably, CasE strongly cleaves the Cse3 recognition hairpin (Supplementary Figure 9), but a single mutation U5A in the Cse3 recognition motif [57] (Cse3*) renders it cleavable only by Cse3 and not CasE. As seen in Figure 3b, the endoRNases’ orthogonality suggests that these set of proteins are usable within the same circuit. Of note, some pairs should be avoided (unless beneficial for circuit design) such as RanCas13b:PguCas13b and CasE:Cse3 (with the wt Case3 recognition site). Given the large number of characterized Cas-family proteins with the ability to recognize and cleave specific RNA recognition motifs, PERSIST has the potential to expand beyond the nine proteins characterized here, making PERSIST scalable towards the construction of large and highly sophisticated genetic circuits.

#### Evaluating composability: cascades and positive feedback

Next we evaluated the composability of the PERSIST platform by layering the endonucleases. In Figure 3c we demonstrate a cascade where one endoRNase (CasE) represses another (Csy4), that in turn represses a reporter. We show reporter EYFP expression as a function of CasE expression (proportional to mKO2 fluorescence) at three different Csy4:CasE ratios and with no Csy4. In the absence of Csy4, EYFP is strongly expressed (gray line). When varying amounts of Csy4 DNA is added and CasE is expressed at low levels, the reporter is repressed strongly. As the level of CasE expression increases, it represses Csy4 and expression of EYFP is restored. Note that as higher ratios of Csy4 to CasE are used, CasE is oversaturated and cannot fully repress Csy4 and therefore full fluorescence levels cannot be restored, likely because Csy4 is a slightly stronger repressor than CasE (see Figure 2). Next, we demonstrate a hybrid endoRNase cascade (Figure 3d). In this cascade CasE activates rather than represses Csy4, which can then, in turn, repress an EYFP reporter. In the absence of Csy4, EYFP is strongly expressed (gray line). When varying amounts of Csy4 DNA is added and CasE is expressed at low levels, Csy4 is not yet activated and the reporter remains highly expressed. As the level of CasE expression increases, it activates Csy4 which represses the reporter.

Importantly, while other post-transcriptional platforms, namely protease-mediated regulation [27, 21, 14], have show repression of another repressor as we have here, none have demonstrated direct activation of a repressor as shown here. Figure 3e shows auto positive feedback where an endoRNase is able to self activate when its recognition site is placed in the 3’ ON-switch of its own transcript. The level of output fluorescence for the positive feedback is similar to the constitutively expressed level.

#### Evaluating modularity: All 16 two-input Boolean logic functions

Finally, we demonstrate the modularity of the platform by combining 5’ OFF and 3’ ON switches on the same transcripts to create all 16 two-input Boolean logic functions (Figure 4). Importantly, multiple recognition sites can be combined within the same 5’ OFF or 3’ ON PERSIST motif (see “NOR” and “OR”) or within separate motifs on the same transcript (see “NIMPLY”). In fact 10 of the 16 logic gates can be constructed with a single transcript and they all perform well with dynamic ranges approaching two orders of magnitude. Overall, these logic gates compare well to previous post-transcriptional platforms as well as DNA-based logic gates in mammalian cells [5, 64, 23].

**Figure 4.**
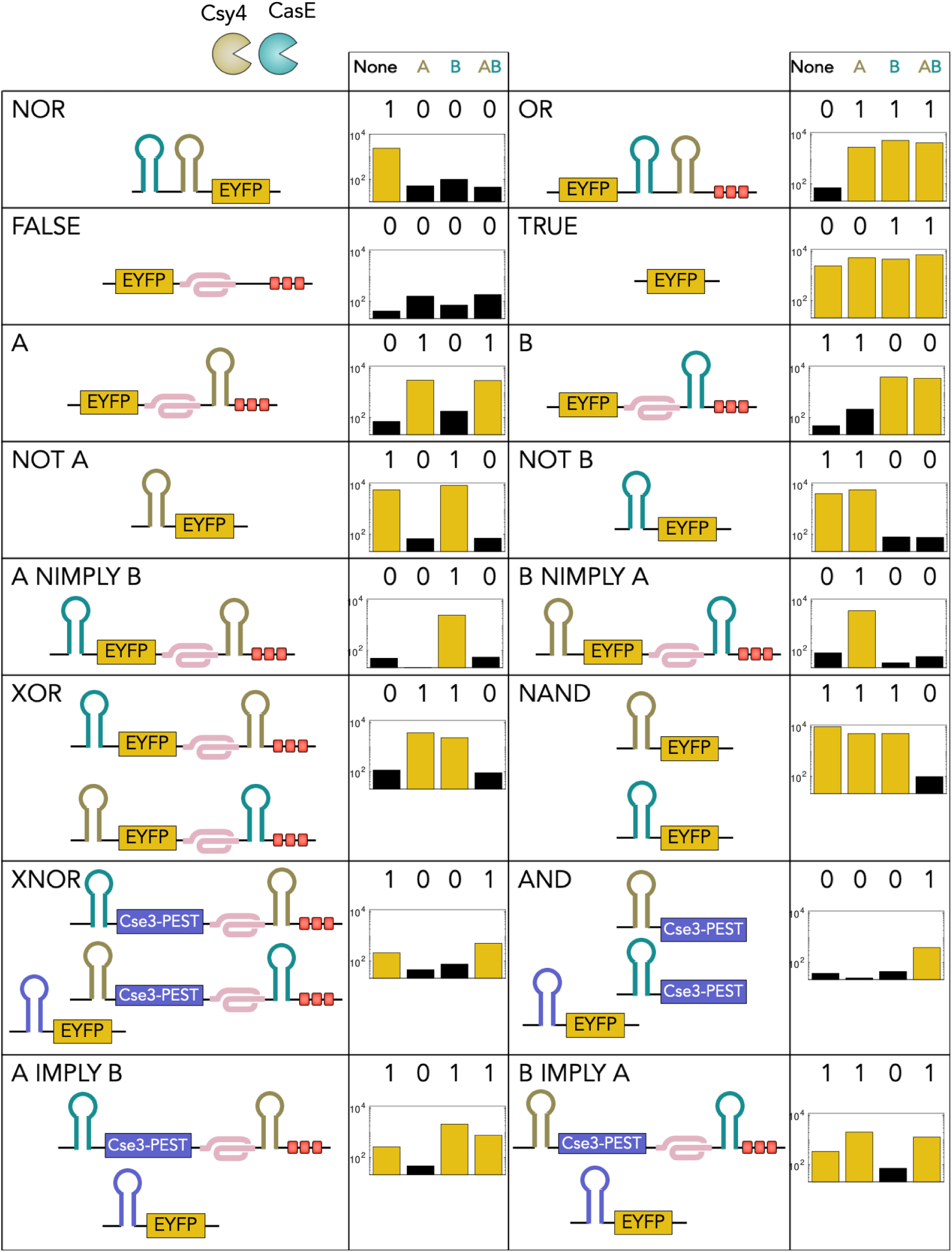
PERSIST endoRNase platform is modular. All 16 two-input Boolean logic functions can be created using Cas endoRNases and the PERSIST ON- and OFF-switches. Each logic function was constructed using EYFP as output with other constructs included as necessary to confer designed response to two inputs: Csy4 and CasE. Cse3 endoRNase was used as an intermediate regulator where necessary. Each set of constructs encoding specific logic functions were evaluated without input endoRNases (‘None’ column), with Csy4 only (‘A’ column), with CasE only (‘B’ column), and with both Csy4 and CasE (‘AB’ column). Expected Boolean logic output is depicted above each chart. Y-axis shows EYFP Fluorescence (a.u.) where each bar represents the median of > 1,000 transfected cells (HEK293FT).

### Dual-function enables compact implementation of complex logic

Importantly, unlike transcription factors in DNA logic-based systems, which usually function to either repress or activate transcription, these endoRNases can act as both “activators” and “repressors” by placing their recognition sequence in a PERSIST-ON or PERSIST-OFF motif (Figure 5a). There are several transcriptional systems that have devised clever ways of switching between activator and repressor [39], but these systems show modest fold change and lack broad composability. In the PERSIST platform, a given endoRNase acts as a repressor if its recognition site is placed in the 5’ PERSIST-OFF motif, while the same endoRNase functions as an activator if its recognition site is placed in the 3’ PERSIST-ON motif. Importanlty, endoRNases can exhibit this dual functionality simultaneously, as shown in Figure 5b, where both a reporter encoding a Csy4-repressible tagBFP transcript and a reporter encoding a Csy4-activatable EYFP transcript can be acted on at the same time in the same cell by Csy4. We took advantage of this feature of the PERSIST platform to compactly build several motifs including a feed-forward loop (FFL) and a single-node positive feedback/repression motif, which in turn enabled a more robust 3-stage cascade and a two-node bistable switch.

**Figure 5.**
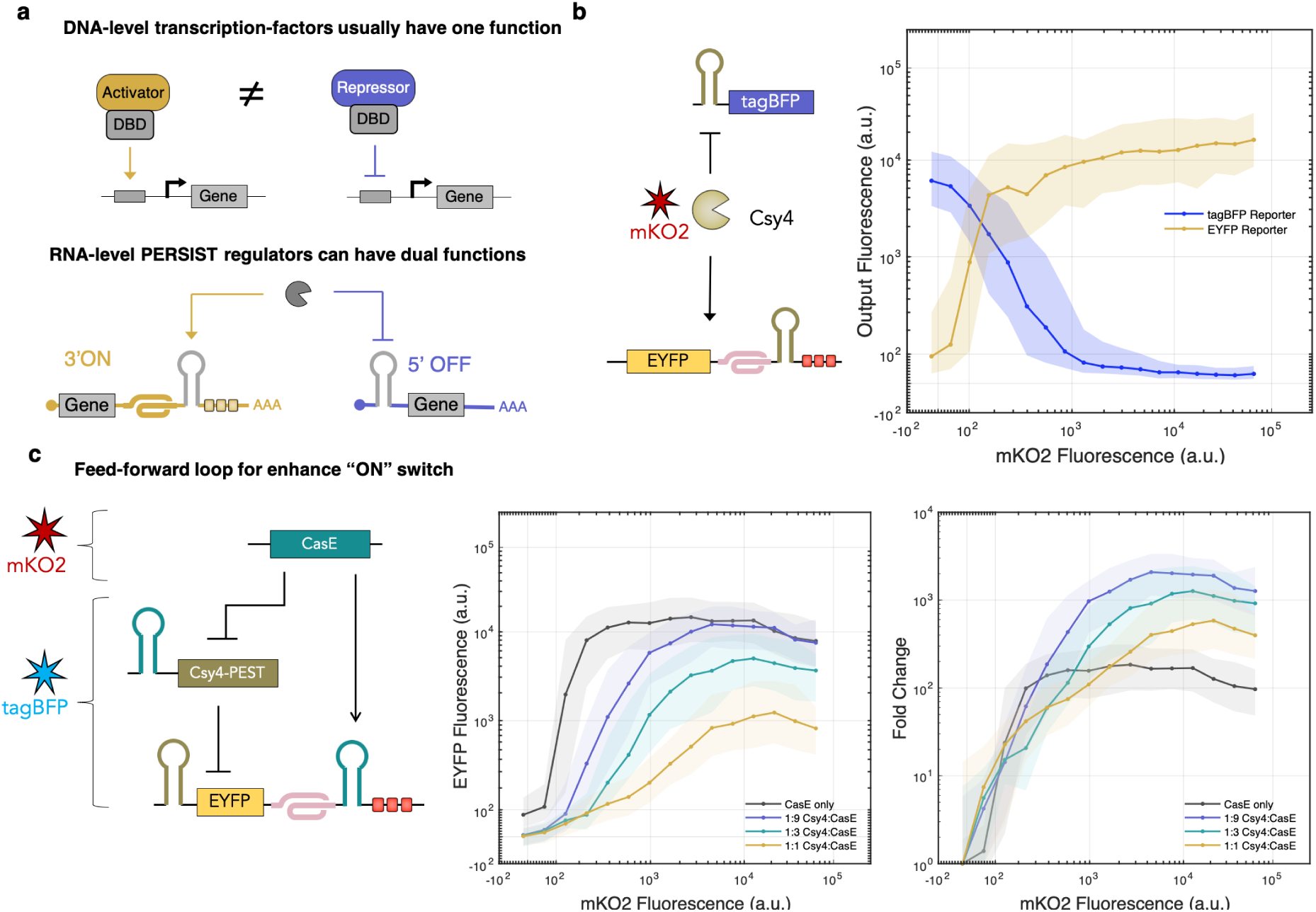
Dual function of Cas endoRNases in the PERSIST platform is advantageous for engineering. **a**, Schematic comparing transcriptional regulation to PERSIST regulation. Transcription factors cannot function as activators and repressors simultaneously while Cas endoRNases can using the PERSIST platform. **b**, A tagBFP-encoding OFF-switch reporter (blue line) and a EYFP-encoding ON-switch reporter (yellow line) can be acted on simultaneously by Csy4 in the same cell. Transfection efficiency of the tagBFP OFF-switch reporter and EYFP ON-switch reporter was tracked by a plasmid encoding constitutive iRFP720 while Csy4 transfection efficiency was tracked by a plasmid encoding constitutive mKO2. Median tagBFP and EYFP (lines) and interquartile range (shaded regions) were evaluated at various mKO2 levels for iRFP720-positive cells. **c**, A coherent feed-forward loop is enabled by dual function of the Cas proteins and improves response of the PERSIST ON-switch motif. For plasmid ratio calculations, CasE plasmid was maintained at 75ng. Transfection efficiency of the EYFP reporter and intermediate Csy4 plasmid was tracked by a plasmid encoding constitutive tagBFP while CasE transfection efficiency was tracked by a plasmid encoding constitutive mKO2. Median EYFP (lines) and interquartile range (shaded regions) were evaluated at various mKO2 levels for tagBFP-positive cells. Fold change (right) was calculated by normalizing to the EYFP value at the lowest mKO2 bin. Note that over 1,000 fold dynamic range is achieved for some Csy:CasE ratios compared to roughly 100 fold dynamic range achieved by the ON-switch alone (gray line). **b,c**, Each sample represents the evaluation of > 1,000 transfected cells (HEK293FT).

#### Feed-forward loop

The coherent feed-forward loop (cFFL) motif provides redundant regulation, where regulation of an output gene via multiple pathways often yields greater dynamic range of downstream gene expression levels. The cFFL is common in many natural biological signalling networks [29]. We hypothesized that a cFFL could increase the dynamic range of the PERSIST-ON response to an endoRNAse input. In our cFFL, CasE acts as the input and Csy4 as an intermediate node, where both act on the output (Figure 5c). When CasE is absent, the reporter is both degraded by the ON switch motif and repressed by Csy4, resulting in much lower background than the PERSIST ON-switch alone. When CasE is present, it directly activates the reporter and indirectly activates it by repressing Csy4. This cFFL is possible because the input endoRNase is able to simultaneously act as an activator for the reporter and as well as a repressor for Csy4, resulting in dynamic ranges of over 1,000 fold and very low background expression (Figure 5c.)

#### Single-node positive feedback + repression

Another interesting motif enabled by the PERSIST device dual function is single-node positive feedback + repression (Figure 6a). We constructed such a motif using Csy4 positive feedback and an EYFP OFF-switch reporter containing the Csy4 recognition site in its 5’ UTR. In a control experiment, when Csy4 functions as just a repressor, it is expressed stably and represses the reporter even at low levels. In a second control experiment, when Csy4 repressor contains the ON switch degradation tags but is unable to self-activate (the ON switch does not contain a recognition site), Csy4 transcript is degraded resulting in poor repression of the reporter. when the Csy4 recognition site is placed in the ON switch motif, Csy4 activates itself through a positive feedback loop and recovers its ability to target and repress the reporter.

**Figure 6.**
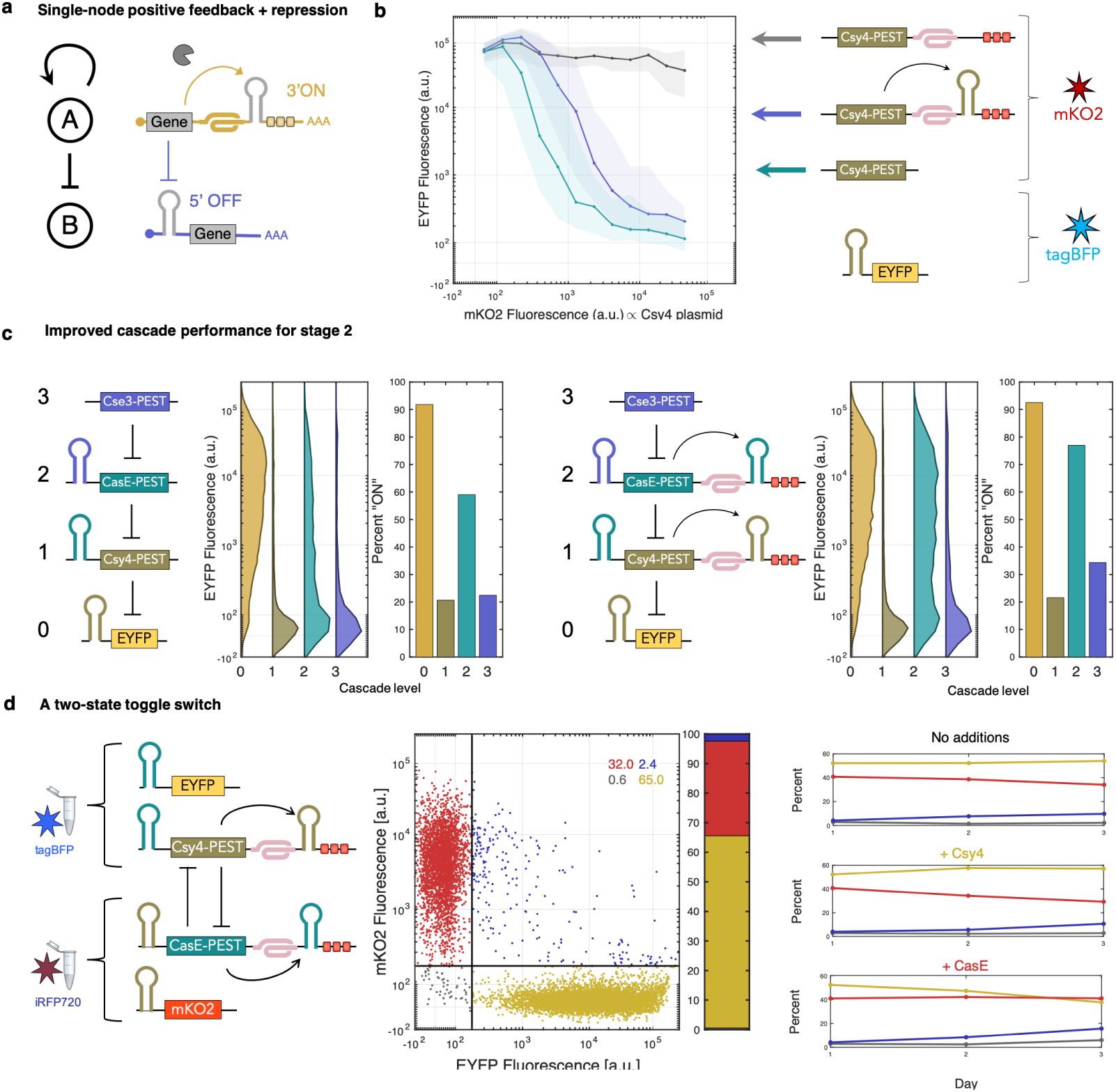
PERSIST dual-function enables single-element positive feedback + repression motif. **a**, Schematic of positive feedback + repression motif. **b**, An endoRNase is able to activate itself through its own PERSIST ON-motif and repress another element, which allows for more graded response. Transfection efficiency of the EYFP reporter was tracked by a plasmid encoding constitutive tagBFP while Csy4-containing plasmid transfection efficiency was tracked by a plasmid encoding constitutive mKO2. Median EYFP (lines) and interquartile range (shaded regions) were evaluated at various mKO2 levels for tagBFP-positive cells. **c**, The positive feedback + repression motif can be used to improve the performance of the second stage of a 3-stage repression cascade. A larger percentage of stage-2 cells exhibit “ON” behavior (right) compared to when repression alone is used with a trade-off existing for third stage response, which increases when positive feedback is used. Each bar represents the evaluation of cells across all transfection levels of each stage. **d**, The positive feedback + repression motif can be used to make a genetic bistable switch. mKO2 and EYFP expression was evaluated for iRFP720- and tagBFP-positive cells. A majority of cells display expression in only a high-mKO2/low-EYFP (red dots) or high-EYFP/low-mKO2 (yellow dots) state. Few cells expressed high levels of both (blue dots) or none (gray dots). Percentages of cells in each state are shown in inset and depicted in bar. Right panel: evaluation of switching in transient transfection. Toggle switch-transfected cells were transfected a day later with inducer endoRNases: either Csy4, CasE, or dummy plasmids and analyzed for two more days. Percentage of cells in each state (high-mKO2/low-EYFP: red lines, high-EYFP/low-mKO2: yellow lines, high-EYFP/high-mKO2: blue lines, and low-EYFP/low-mKO2: gray lines) were calculated as in dot plot to left. Inducer endoRNase transfection efficiency was not tracked with fluorescent proteins so values represent evaluation of all cells regardless of transfection state. Data shows that a larger percentage of cells transfected with endoRNase show switching to the expected state compared to a toggle control sample where no inducer endoRNase was introduced. **b,c,d**, Each sample represents the evaluation of > 1,000 transfected cells (HEK293FT).

We further demonstrate the robustness of this motif through the construction of a 3-stage repression cascade (Figure 6c). In the repression-only cascade, there is indeed a drop in signal restoration at stage 2 (Figure 6c, left), however this cascade still outperforms previous RNA-based repression cascades. When positive feedback is included, a larger percentage of stage-2 cells exhibit “ON” state behavior (Figure 6c, right). There is, however, a trade off as a larger percentage of stage 3 cells exhibit incorrect “ON” behavior when positive feedback is included.

#### Construction of an endoRNase-based bistable switch

To demonstrate this motif’s utility we sought to create a bistable toggle switch. Notably, it has already been shown that DNA-based switch topologies that include positive feedback loops at each node improve bistability compared to those with cross-repression alone [45] and similarly that RNA-based switches relying only on cross-repression do not exhibit bistability in an uninduced state [66]. Here, our positive feedback + repression motif enables compact construction of a bistable toggle switch that requires only two nodes encoding both cross-repression and positive feedback. Csy4 and CasE both activate themselves as well as repress the other endoRNase (Figure 6d). We design separate constructs containing fluorescent proteins (mKO2 and EYFP) that can be repressed by each of the endoRNases to track cell state. We then evaluated these constructs via transient poly-transfection [26] to allow for many ratios of each node to be sampled (Figure 6d, middle). We achieved thorough sampling of node ratios by creating separate poly-transfection complexes for each node; we tracked the transfection efficiency of the Csy4 node containing the endoRNase plasmid and EYFP output using a constitutive tagBFP reporter while we measured the transfection efficiency of the CasE node containing the endoRNase plasmid and mKO2 output using an iRFP720 reporter. Cells transfected with both nodes (determined by tagBFP and iRFP720 fluorescence) exhibited two major states: a high-mKO2/low-EYFP state and low-mKO2/high-EYFP state. Promisingly, despite the wide range of ratios sampled, 97% of cells were in one state or the other and only 3% of cells expressed none or both fluorescent proteins indicating that this system is indeed bistable. Next, we show that it is possible for the system to be switched between states. The same poly-transfection as described above was performed and after one day (once the system had already assumed bistability) we transfected plasmids encoding either Csy4 or CasE. Despite the system having already segregated into two mutually exclusive states cells were still able to switch states once transfected with either endoRNase in the switch: when cells were transfected with Csy4, an increased percentage of cells switched to the low-mKO2/high-EYFP state, while when cells were transfected with CasE there was a higher percentage of cells in the high-mKO2/low-EYFP state. This switch could serve as a useful tool for studying cell fate decision making, development of cell type classifiers, and the creation of gene and cell therapies where strict ON or OFF states are required instead of a graded response, for example in CAR T-cell activation or kill switches.

## DISCUSSION

Here we present a fully composable, scalable, and programmable RNA activation/repression platform. Importantly, we demonstrate that post-transcriptional regulation with the PERSIST platform enables the use of constitutive promoters that resist epigenetic silencing. In particular, development of the RNA-level ON switch, which has been a challenge in the field due to the fact that translation is typically “on” by default, is especially enabling as it allows for direct positive regulation by a range of cleavage effectors. We found that CRISPR endonucleases exhibit roughly 100 fold dynamic range in repression and roughly 10 fold dynamic range in activation through the PERSIST-OFF and -ON switches respectively. Using these endoRNases as RNA-level activators and repressors we demonstrate a range of genetic computations that give us confidence that this system could be used in place of transcription factor-based regulation in scenarios where epigenetic silencing could be problematic. Unlike transcriptional logic which often involves optimizing various repeats of operons with careful placement with respect to minimal promoters to achieve the desired response, programming with the PERSIST platform is straightforward and involves placing the target recognition sites either in the 5’ UTR PERSIST-OFF position or the 3’ UTR PERSIST-ON position. In fact, CasE, one of the most robust endoRNases explored here has already been used to implement an incoherent feedforward loop that achieves control of gene expression regardless of DNA copy number and resource competition [37]. In particular, we also demonstrate that the unique dual functionality of these endoRNases in the PERSIST platform enables facile engineering of useful motifs. These motifs allowed for improved signal restoration and the creation of a compact RNA-level bistable toggle switch. Taken together, the robustness, scalability, and modularity of the platform and its ability to resist silencing promises the programming of more sophisticated and reliable behavior in gene and cell therapies.

## METHODS AND MATERIALS

### Plasmid cloning

Hierarchical Golden Gate Assembly was used to assemble all expression constructs from “Level 0” (pL0) parts as published previously [17]. Here we create sub-pL0s which contain the PERSIST ON-switch stabilizer, cut site and degradation domains, which can be assembled using BbsI golden gate into a 3’ UTR pL0. We also devise a “Level 2” cloning scheme using SapI golden gate reactions to assemble multiple transcription units onto a single plasmid.

### Cell culture

HEK293FT (Invitrogen) cells were maintained in Dulbecco’s modified Eagle medium (DMEM, Corning) supplemented with 10% FBS (Corning), 1% Penicillin-Streptomycin-L-Glutamine (Corning) and 1% MEM Non-Essential Amino Acids (Gibco). CHO-K1 cells (ATCC) were maintained in Ham’s F-12K (Kaighn’s) medium supplemented with with 10% FBS (Corning), 1% Penicillin-Streptomycin-L-Glutamine (Corning), and 1% MEM Non-Essential Amino Acids (Gibco). Doxycycline hyclate (Dox, Sigma-Aldrich) was diluted in water at 10mg/uL, stored at −20C, and used at a final concentration of 4uM for experiments. Trichostatin A (TSA, Sigma-Aldrich) was diluted in DMSO, stored at −20C and used at a final concentration of 100nM for experiments.

### Transfections

HEK293FT cell transfections were performed using Lipofectamine 3000 (invitrogen) with a 1.1:1.1:1 Lipofectamine 3000:P3000:DNA ratio in 24-well or 96-well format. Complexes were prepared in Opti-MEM (Gibco). Cells were plated on the day of transfection in culture media without antibiotics and analyzed by flow cytometry after 48 hours. CHO-K1 cell transfections were performed using Lipofectamine LTX (invitrogen) with a 4:1:1 LTX:Plus Reagent:DNA ratio in 24-well or 96-well format. Complexes were prepared in Opti-MEM (Gibco). Cells were plated on the day of transfection and analyzed by flow cytometry after 48 hours.

### Genomic Integration

Constructs were integrated into CHO-K1 cells that had been previously engineered with a landing pad in their putative Rosa26 locus [24]. For payload integration, cells were transfected with engineered construct and BxBI expressing plasmid in 12-well plate format. After 3 days, media was supplemented with 8ug/mL puromycin (invivogen) and 10ug/mL blasticidin (invivogen). Cells were maintained under selection for two weeks with fresh media containing antibiotics refreshed every 2 days. After two weeks, cell lines were evaluated for response and underwent single-cell sorting.

### Flow Cytometry and single-cell sorting

Cell fluorescence was analyzed with LSR Fortessa flow cytometer, equipped with 405, 488, 561, and 637 nm lasers (BD Biosciences). The following laser and filter combinations were used to evaluate the fluorescent proteins used in transient transfection experiments this study: tagBFP, 405nm laser, 450/50 filter; EYFP, 488nm laser and 515/20nm filter; mKO2, 561nm laser, 582/42 filter; iRFP, 640nm laser, 780/60 filter. Cell sorting was performed on a FACSAria cell sorter, equipped with 405, 488, 561 and 640nm lasers. The following laser and filter combinations were used to evaluate the fluorescent proteins used in single-cell sorting: EBFP, 405nm laser, 450/40 filter; EYFP, 488nm laser, 530/30 filter; mKO2, 561nm laser, 582/15 filter.

## Data Analysis

Analysis of flow cytometry data was analyzed in MATLAB with a custom analysis pipeline, which will be made available. For each data set, a compensation matrix is computed from single color controls to account for spectral bleed-through. After compensation, thresholds were set to evaluate positively transfected cells. For single construct analyses, linear fits were computed between EYFP and tagBFP expression levels. For poly-transfection involving multiple plasmids, data was binned before linear fits or summary statistics were calculated.

## ACKNOWLEDGMENTS

We thank Kevin Lebo and Ross Jones (MIT) for discussion, Jin Huh (MIT) for parts, Leonid Gaidukov for providing landing pad cells, Allen Tseng for BxbI plasmid, Emerson Glassey for SapI cloning overhang analysis, and the Koch Flow Cytometry Core for assistance with single cell sorting.

