## Supplemental Information for "PERSIST: A programmable RNA regulation platform using CRISPR endoRNases"

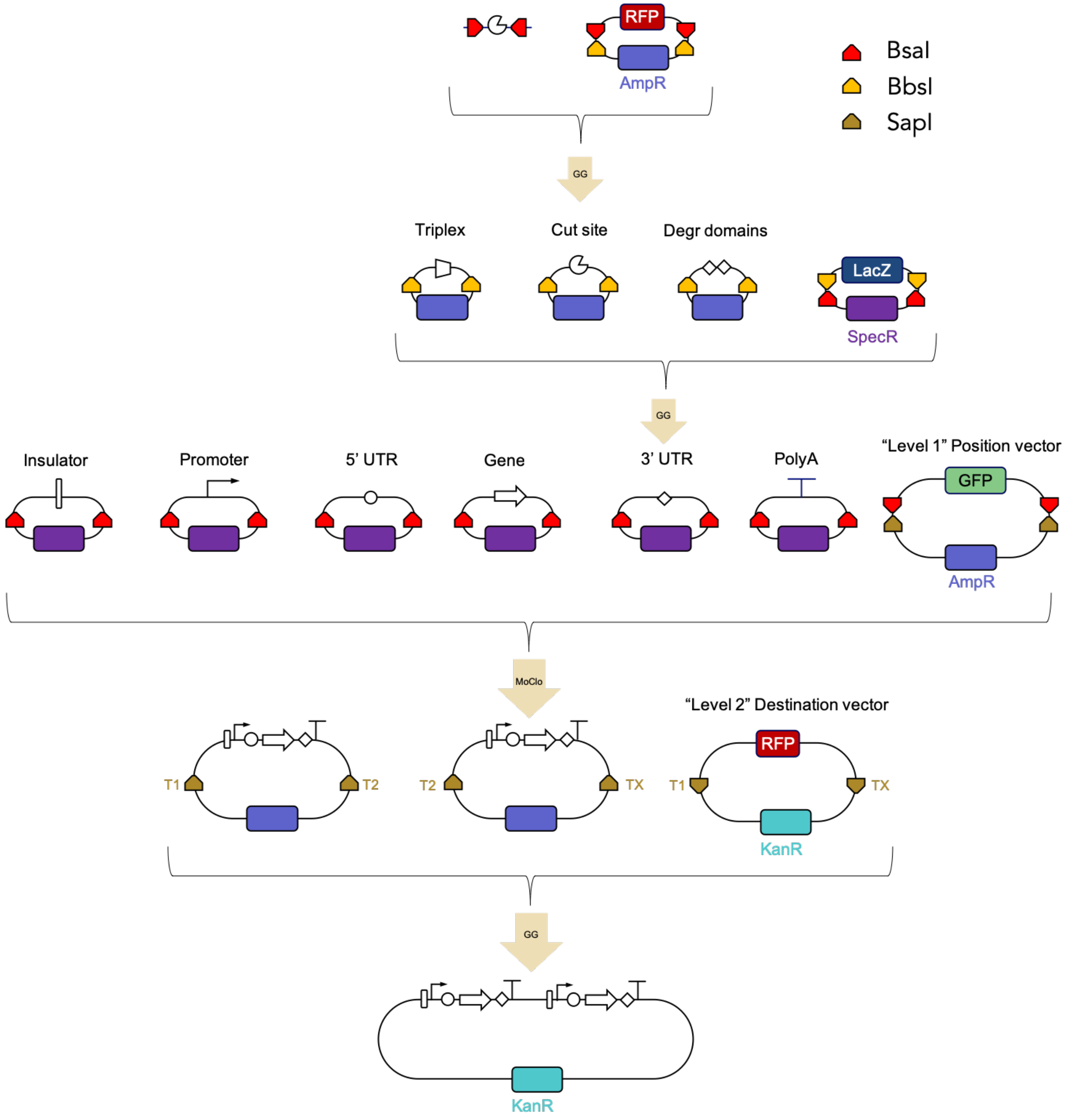


**Supplementary Figure 1. Hierarchical golden gate cloning method.** sub-pL0s are created using BsaI golden gate reactions into sub-pL0 backbones containing ampicillin resistance gene. pL0[3'-UTR] position vectors can be made through BbsI golden gate reactions with sub-pL0 vectors and pL0[3'-UTR] backbone vector containing a gene for spectinomycin resistance. pL1 expression vectors can be made through BsaI golden gate reactions with pL0 vectors and pL1 position vector backbone vector containing a gene for ampicillin resistance. Finally, multi-transcription unit pL2s are created through SapI golden gate reactions with pL1 vectors and pL2 backbone vector containing a gene for kanamycin resistance.

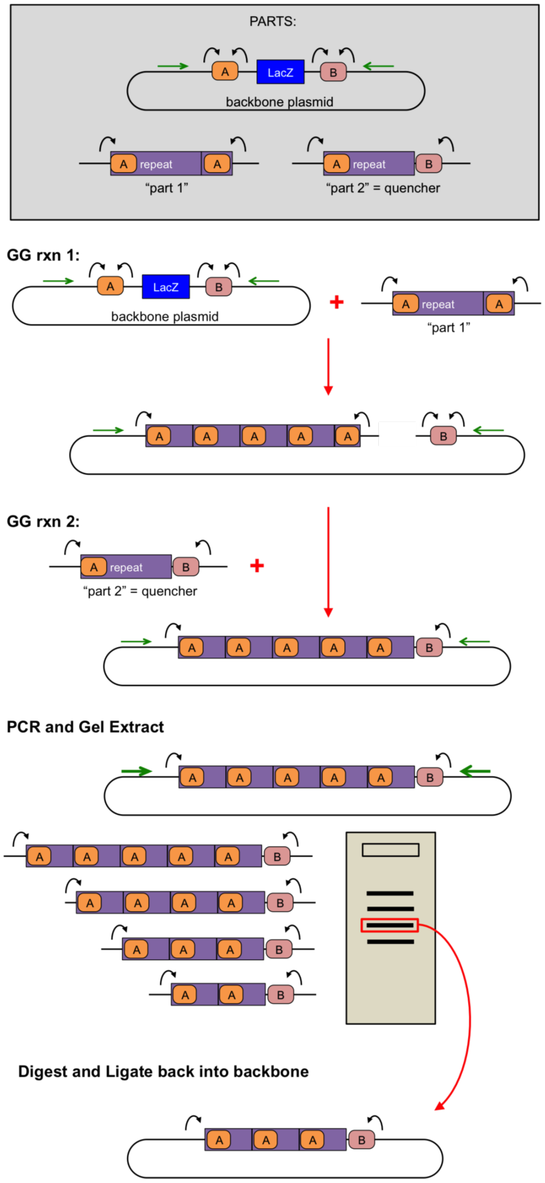


**Supplementary Figure 2. "NxGG": In-house strategy for developing constructs with many sequence repeats.** Briefly, the protocol works by designing a repeat element containing Type IIs restriction sites with the same overhangs on both sides ("part 1"). A GG reaction step with the backbone will allow part 1 to concatenate on the backbone. Adding the quencher, "part2", binds to part1 and the backbone to close the plasmid and complete the reaction. When directly transformed the reaction has significant bias towards only one repeat so generally it is best to PCR from the backbone of the plasmid, gel extract the band of choice with selected number of repeats and re-digest and ligate back into the backbone.

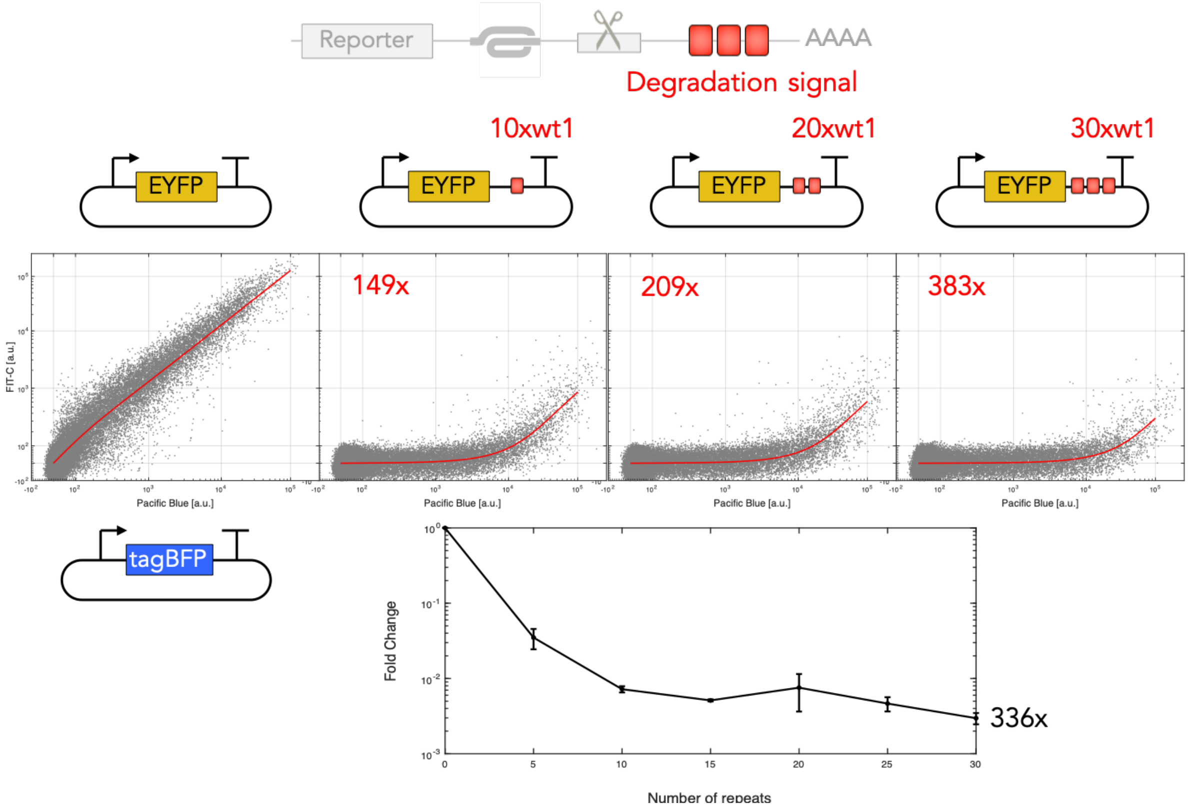


**Supplementary Figure 3. Data analysis for 1D co-transfection.** We assume that transcript degradation machinery is not limiting and thus compute linear fits of the reporter and transfection marker fluorescence. This results in a holistic evaluation of transfection data which typically takes into account $>$ 10,000 cells and does not bias analysis based on transfection marker bin choice. This same method was also used to evaluate ribozyme response in cis as in Figure 2b.

**Analyzing trans-response for PERSIST OFF-switches by 2D poly-transfection**

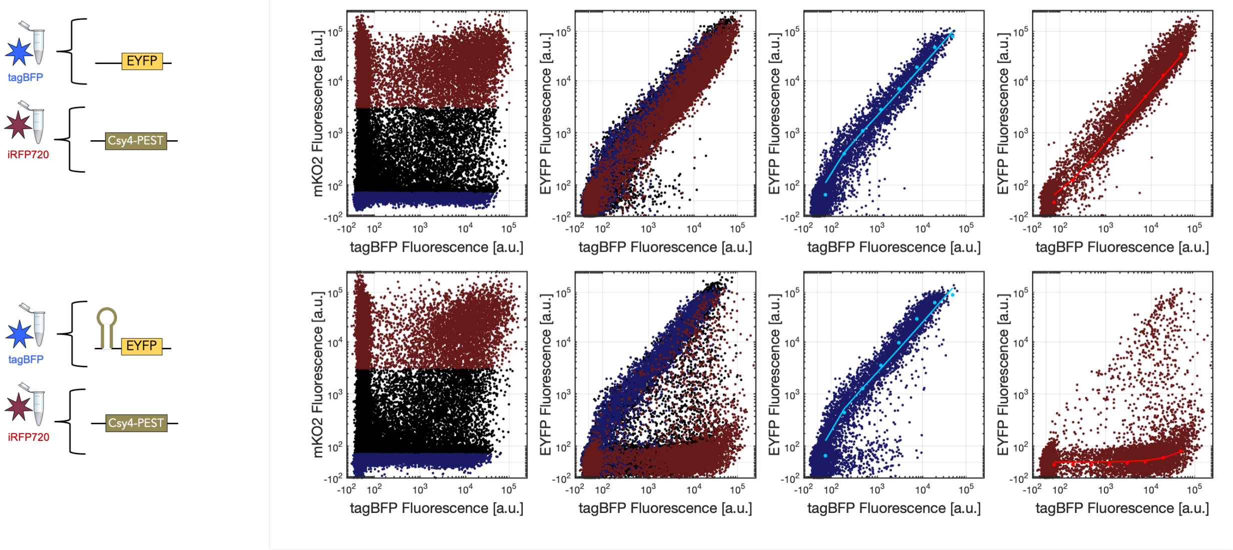


**Analyzing trans-response for PERSIST OFF-switches by 2D poly-transfection**

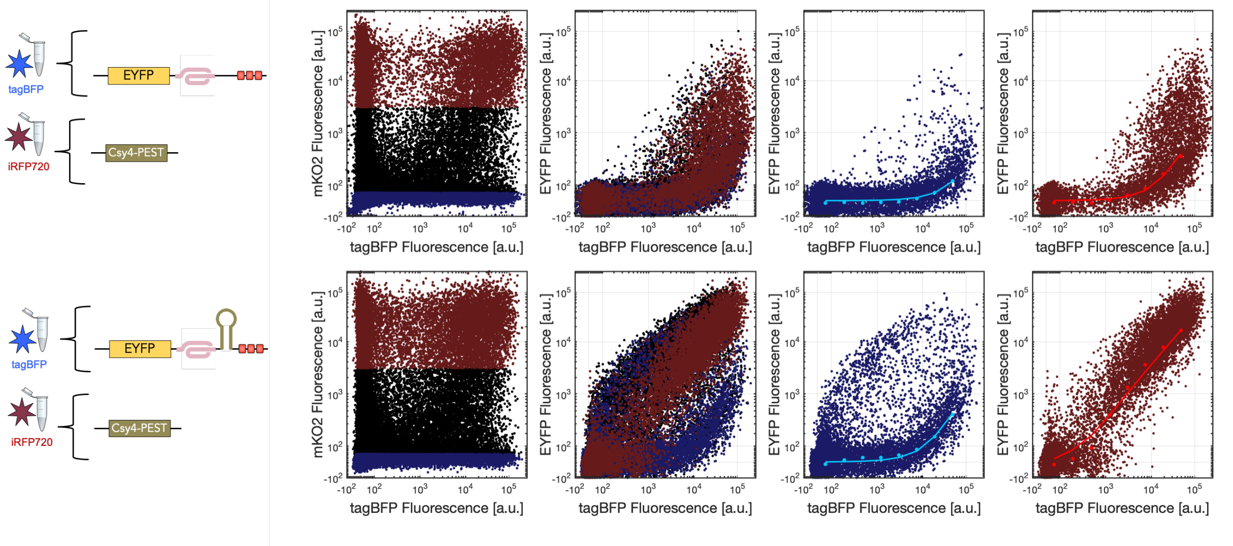


**Supplementary Figure 4. Analysis of endoRNase PERSIST platform responses using 2D poly-transfection** Here we demonstrate the evaluation of Csy4 in the PERSIST OFF- and PERSIST ON-switches. For either OFF- or ON-switch analysis of Csy4 response, switch constructs were compared to similar control constructs that did not contain the Csy4 recognition site. Csy4 plasmid transfection efficiency was tracked by a plasmid encoding constitutive mKO2 while EYFP reporter plasmid transfection efficiencies were tracked by a plasmid encoding constitutive tagBFP. The first column shows the poly-transfection space for each reporter with Csy4 which are similar across reporters and sample the 2D transfection space well. Data was then categorized as either having low (blue dots) or high (red dots) mKO2 fluorescence which is proportional to Csy4 plasmid transfection efficiency. We see that at high Csy4 values where the recognition site is present, we see an expected shift in population. Each of these populations were binned on tagBFP fluorescence and summary values were fit to an extreme value distribtion using the $evfit$ function in MATLAB. Linear fits of these summary values could then be fit as in the 1D calculations. Fold change values were then computed as (sample slope at high mKO2/sample slope at low mKO2)/(control slope at high mKO2/control slope at low mKO2). This method therefore takes into account and normalizes by any non-specific effects observed by added endoRNase alone. This same method was also used to evaluate miRNA response as in Figure 2c

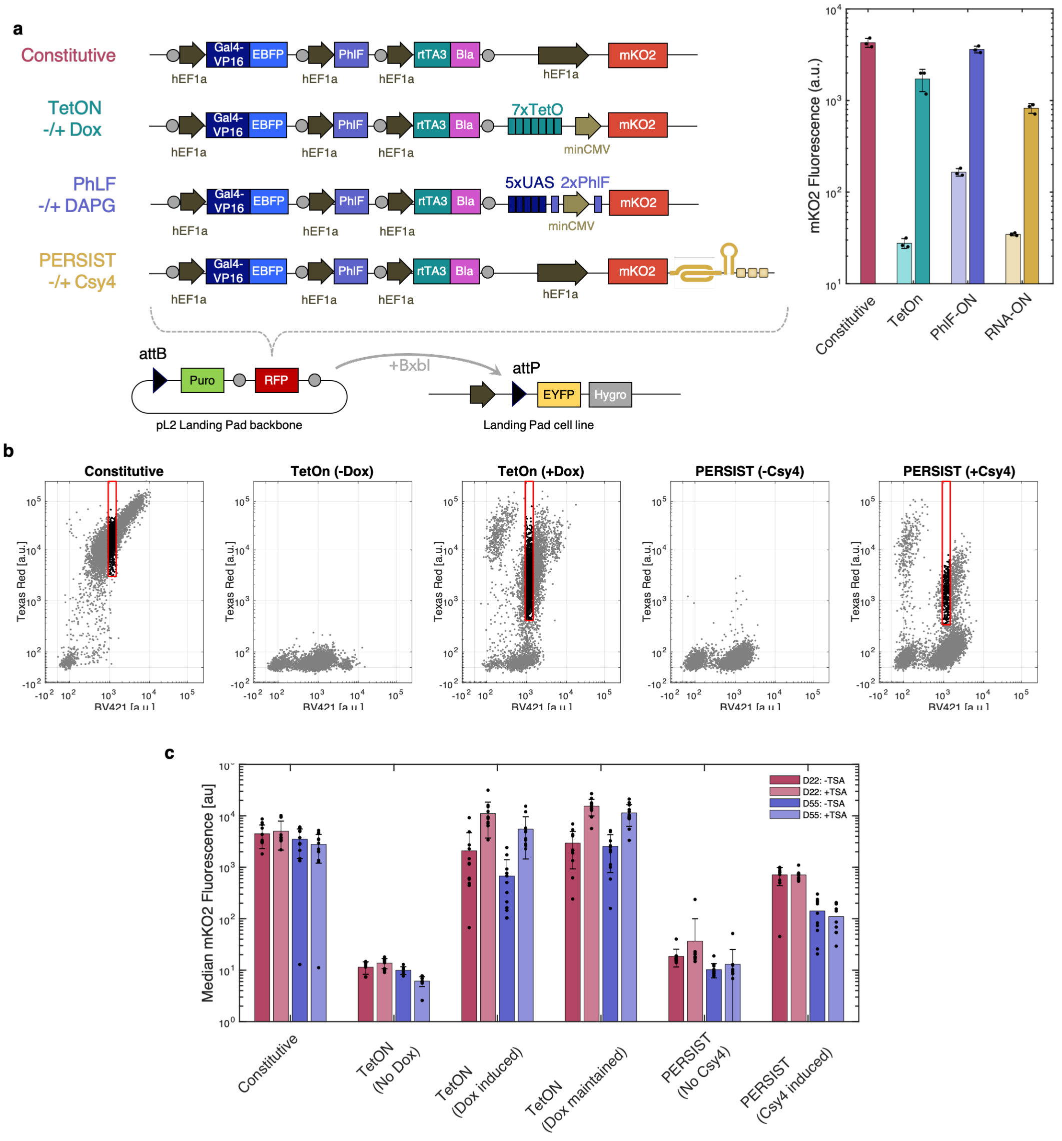


**Supplementary Figure 5. Generating cell lines to evaluate silencing. a,** Circuits for each switch were assembled into a pL2 vector containing necessary attB sites for landing pad integration. Parts for all circuits were cloned upstream of each switch in order to isolate any differences to the last transcription unit to compare each switch. Plasmids were integrated into Rosa26 LP CHO-K1 cell lines using BxbI and under puromycin and blasticidin selection populations were seen that exhibited a shift from EYFP to EBFP implyin ghte circuit was integrated successfully. Poly-clonally integrated lines were induced and evaluated for their response (right). The PhlF-On circuit did not behave as expected due to high background so was not continued for single-cell sorting. **b,** For single-cell sorting, the cell lines were induced with either Dox or Csy4 and compared to lines that were uninduced to evaluate threshold for sorting. a 96-well plate of each cell line was collected, except for the Tet-On line where two plates were collected for which one was maintained in 4uM Dox and the other was not. The following EBFP gate was used to sort all cell lines: Blue $>$ 925 and $<$ 1432. Importantly, cells were sorted only if they exhibited high response to induction to ensure a positive baseline for all samples. The following mKO2 threshold were used for each circuit: Constitutive $>$ 2922, TetOn $>$ 445, PERSIST $>$ 373. **c,** 12 single-cell-sorted lines were selected randomly for further culturing and response evaluation. Cell lines were induced with either Dox or Csy4 and at the same time were evaluated in the presence or absence of 100nM TSA. Dots show median mKO2 fluorescence for cell lines that had at least 5 cells and bars represent mean ± s.d. of these 12 biological replicates. As shown both the Constitutive control circuit and persist show little change in response to TSA addition, however the Tet-On system response to Dox is increased about 10-fold in the presence of TSA, implying that the cells are experiencing epigenetic silencing.

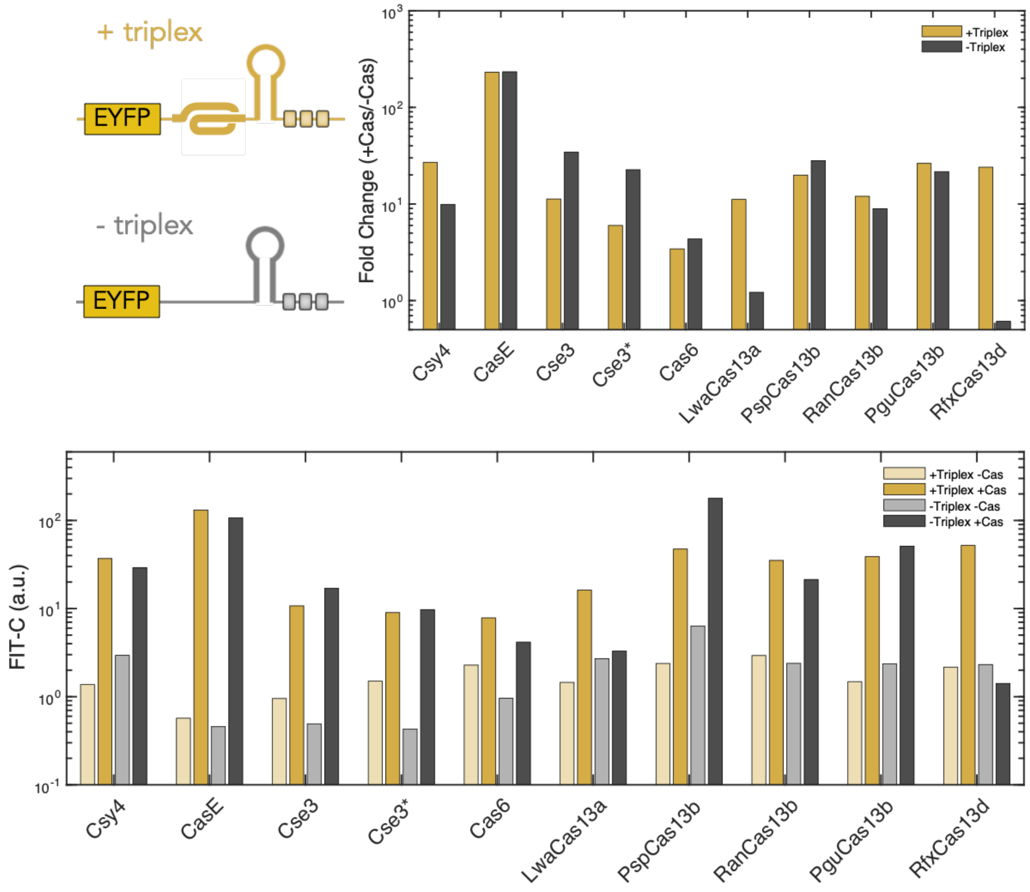


**Supplementary Figure 6.** Evaluating triplex requirement in PERSIST ON switch. Reporters were made for each endoRNase that did or did not contain the MALAT1 triplex. Slopes were calculated as in Supplementary Figure 4. Fold change between +/- endoRNase is reported in top plot. For some endoRNases (e.g. Cse3), removing the triplex resulted in a decrease in fluorescence background while recovery to the same level could still be accomplished resulting in a larger possible dynamic range. Not that for endoRNases that remained bound to the 3' cleaved product (i.e. LwaCas13a and RfxCas13d) the triplex was required for transcript stabilization post-cleavage. Optimal constructs were chosen for each endoRNase.

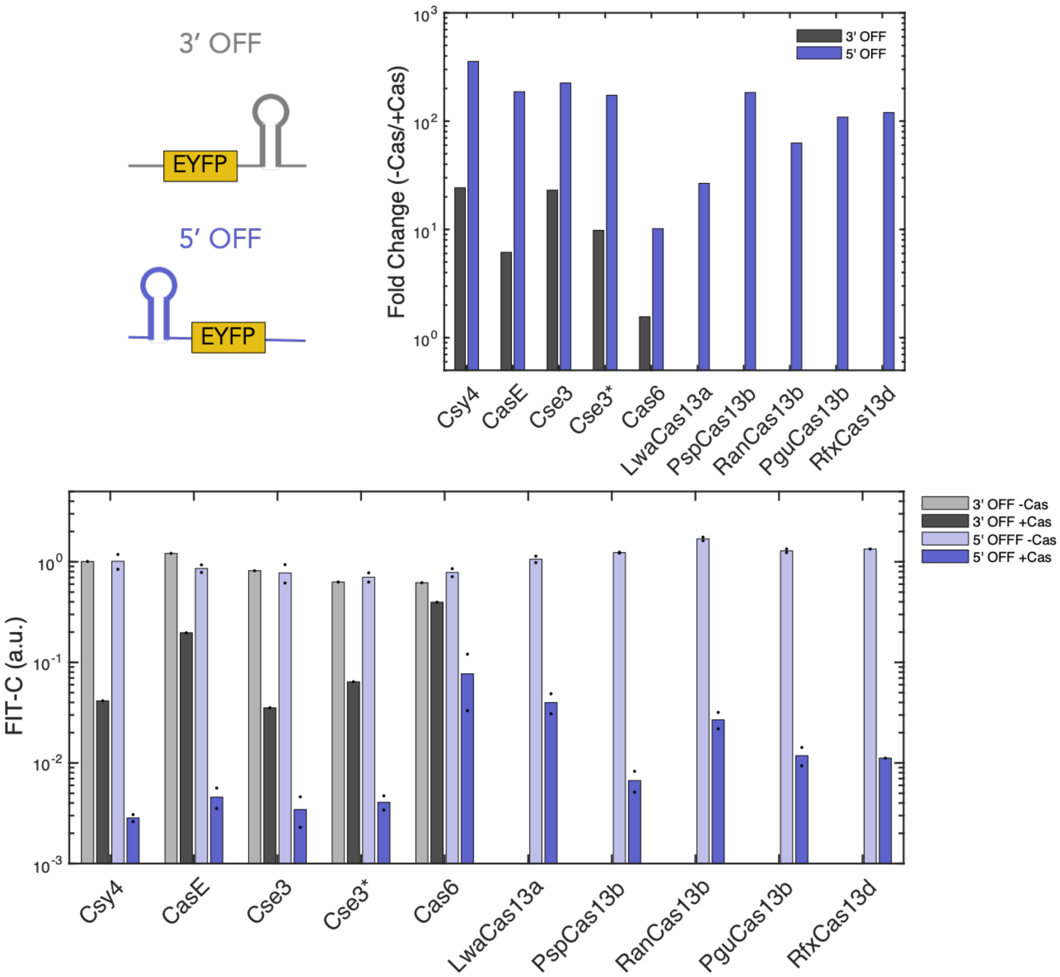


**Supplementary Figure 7.** Evaluating PERSIST OFF switch. EndoRNase response was evaluated and computed as in Supplementary Figure 4 for each construct. For the Cas6 family of endoRNases cleavage sites were placed in either the 5’ or 3’ UTR to evaluate performance. For all cases, placing the endoRNase recognition site in the 5’ UTR resulted in improved repression of expression in the presence of endoRNase. This is likely because these endoRNases stay bound to the 5’ cleaved product and would potentially protect the transcript from degradation in the 3’ UTR as observed in the samples without triplex in Supplementary Figure 6. All endoRNases evaluated here performed well with dynamic ranges of over 10-fold for all endoRNases with the majority approaching 100-fold. The range of repression extents could prove useful in programming and tuning specific circuit behavior.

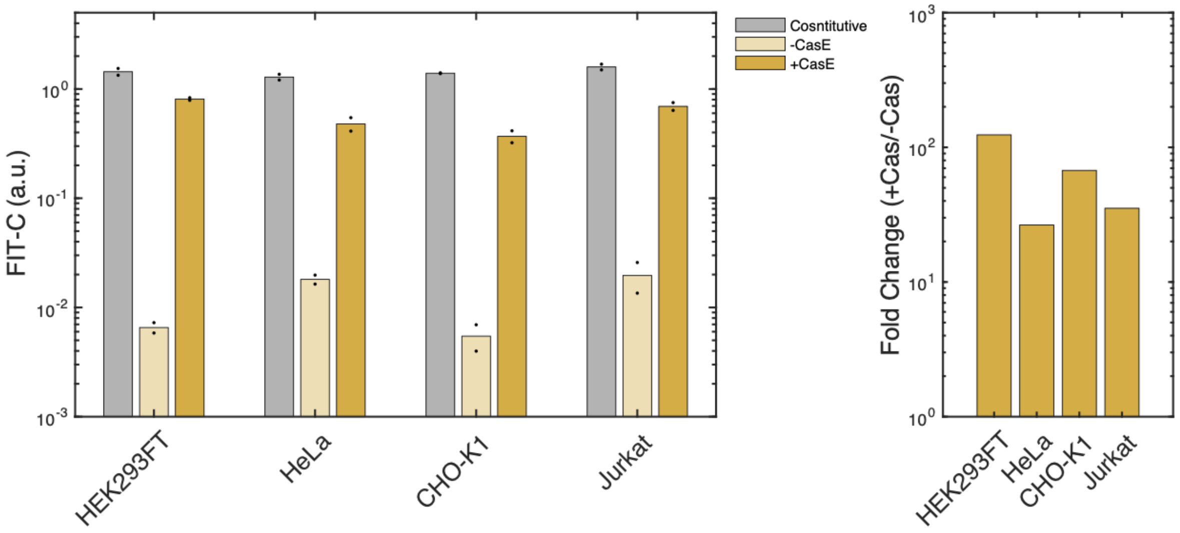


**Supplementary Figure 8.** PERSIST platform functions in a variety of cell types. The CasE-responsive PERSIST-ON switch EYFP reporter was transiently transfected and evaluated in four different cell types. All switches perform well where the switch is degraded effectively in the absence of CasE but approaches constitutive EYFP expression levels in the presence of CasE.

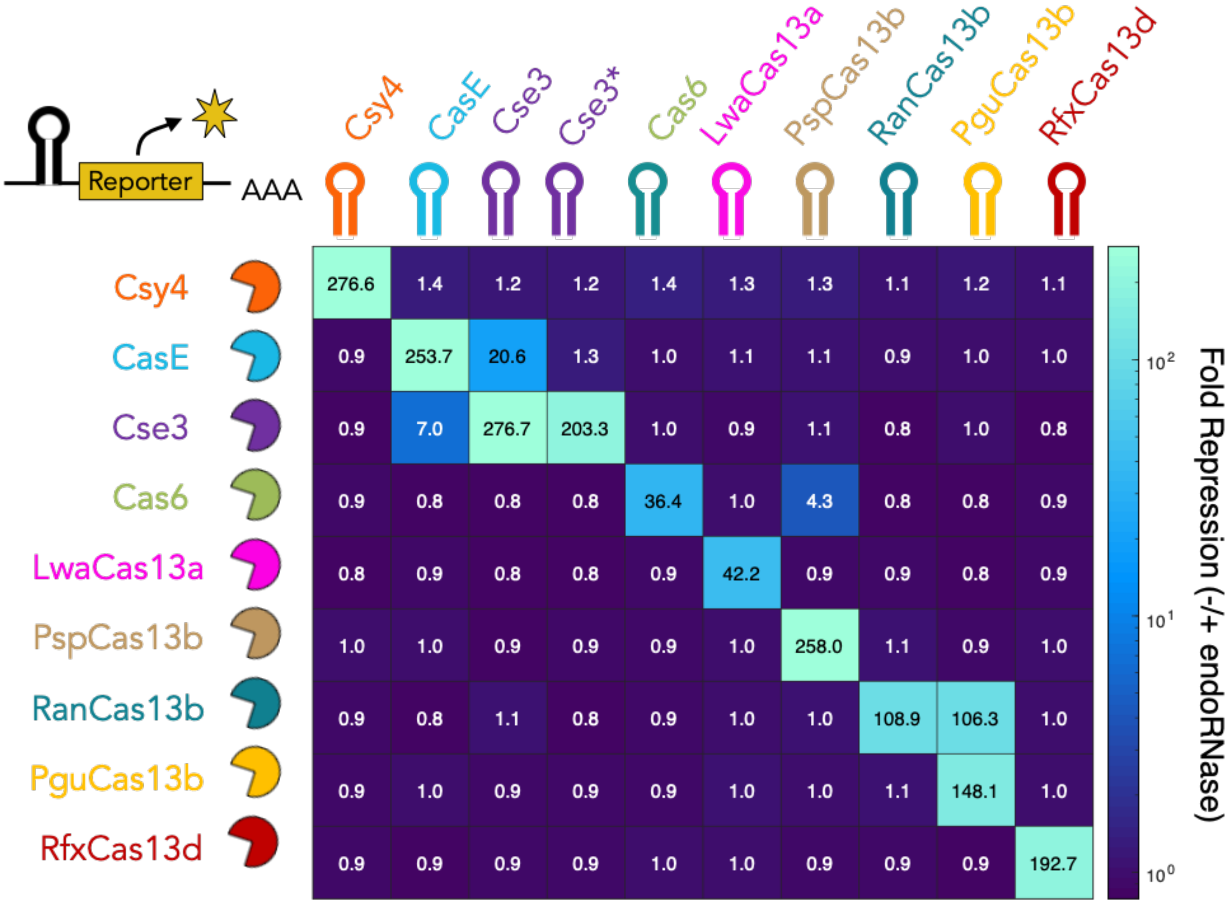


**Supplementary Figure 9. Full orthogonality matrix** Orthogonality of all endoRNase: PERSIST-OFF reporter pairs were evaluated via poly-transfection. Fold responses were computed as in Supplementary Figure 4 compared to a control reporter with no cleavage site. the computed values are shown in each square. Here we see the overlap in cleavage between Cse3 and CasE which can be minimized using the mutated Cse3 hairpin.
